# Moving beyond *P* values: Everyday data analysis with estimation plots

**DOI:** 10.1101/377978

**Authors:** Joses Ho, Tayfun Tumkaya, Sameer Aryal, Hyungwon Choi, Adam Claridge-Chang

## Abstract

Over the past 75 years, a number of statisticians have advised that the data-analysis method known as null-hypothesis significance testing (NHST) should be deprecated (Berkson, 1942; Halsey et al., 2015; Wasserstein et al., 2019). The limitations of NHST have been extensively discussed, with a broad consensus that current statistical practice in the biological sciences needs reform. However, there is less agreement on reform’s specific nature, with vigorous debate surrounding what would constitute a suitable alternative (Altman et al., 2000; Benjamin et al., 2017; Cumming and Calin-Jageman, 2016). An emerging view is that a more complete analytic technique would use statistical graphics to estimate effect sizes and evaluate their uncertainty (Cohen, 1994; Cumming and Calin-Jageman, 2016). As these estimation methods require only minimal statistical retraining, they have great potential to shift the current data-analysis culture away from dichotomous thinking towards quantitative reasoning (Claridge-Chang and Assam, 2016). The evolution of statistics has been inextricably linked to the development of quantitative displays that support complex visual reasoning (Tufte, 2001). We consider that the graphic we describe here as *estimation plot* is the most intuitive way to display the complete statistical information about experimental data sets. However, a major obstacle to adopting estimation plots is accessibility to suitable software. To lower this hurdle, we have developed free software that makes high-quality estimation plotting available to all. Here, we explain the rationale for estimation plots by contrasting them with conventional charts used to display data with NHST results, and describe how the use of these graphs affords five major analytical advantages.

## Introduction

### The two-groups design is fundamental

While NHST limits the analyst to the ill-conceived question of ‘Does it?’ (McCloskey, 2002), estimation instead draws the analyst’s attention to the question of ‘How much?’ — the very topic that defines quantitative research. A fundamental research tool is an experiment that uses control and intervention samples: the *two-groups* design. Two-groups data are traditionally analyzed by Student’s *t*-test and its variants. We will use a series of visualizations of two-groups data to illustrate the progression from NHST to estimation-oriented analysis.

### Significance tests obscure two data aspects

The Student’s *t*-test makes the assumption that two groups have identical means (i.e. it proposes that the effect size is zero). It then challenges this null hypothesis with the observed data, by calculating the chance of seeing the observed effect size (or greater) within the hypothesized null distribution—this is the *P* value. If the probability falls below a certain threshold (typically *P* < 0.05), the null hypothesis is rejected. The analyst then plots the two groups’ means in a bar chart and denotes ‘significance’ by marking it with a star (Figure 1A). This visualization has two important deficiencies. First, by displaying only the means and width of their errors, a bar chart obscures the observed values (2014). Second, NHST plots show only the test result (as indicated by a star or a *P* value), while omitting a diagram of the null distribution itself. The omission of the full dataset and distributional information in *t*-tests is a reflection of how NHST—by focusing on an accept/reject dichotomy—diverts attention from effect quantification.

**Figure 1.**
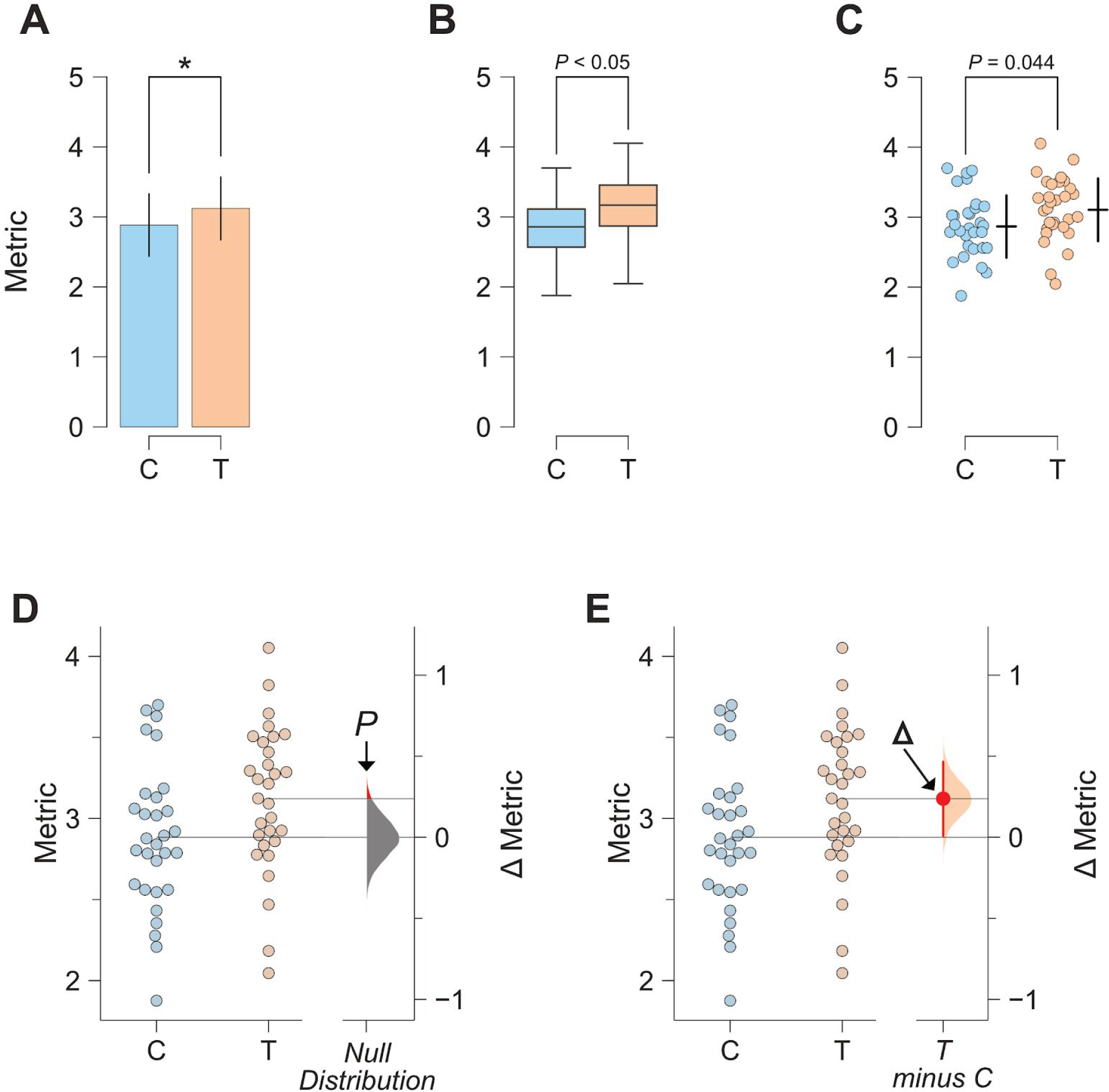
The evolution of two-groups data graphics. **A**. Two-groups data presented in a bar chart. Control (C) and test groups (T) are shown as blue and orange bars, respectively. **B.** The same data presented as a box plot. **C.** A scatter plot (with jitter) allows for all observed values to be visualized (alongside appropriately-offset crossbars to indicate the mean and standard deviation for each group), but does not illustrate the groups’ comparison (ie. the effect size). **D.** A visualization of the two-groups comparison from the null-hypothesis significance testing perspective. The filled curve on the difference axis indicates the null-hypothesis distribution of the mean difference. By definition, this distribution has a mean difference of zero. The area of the red segment indicates the *P* value (for one-sided testing). **E.** An estimation plot uses the difference axis to display on an effect size, here the mean difference (∆). The filled curve indicates the resampled ∆ distribution, given the observed data. Horizontally aligned with the mean of the test group, the ∆ is indicated by the red circle. The 95% confidence interval of ∆ is illustrated by the red vertical line.

The transparency of bar charts is only modestly improved with box plots (Figure 1B); although they do outline more distributional information, box plots do not display complex attributes (e.g. bimodality) or the individual values (Matejka and Fitzmaurice, 2017).

Data-transparency is facilitated with the use of dot plots that show every datum (Cleveland and McGill, 1984; 2017) (Figure 1C). Dot plots are best drawn as beeswarm plots, which convey histogram-like information about the distribution while still displaying every observation (Ecklund, 2015; Waskom et al., 2016; Wilkinson, 1999) (Figure 1D).

### Even when fully visualized, significance tests are misleading

An NHST plot can be made more informative by including a second axis to the side of the observed values (Gardner and Altman, 1986). This *difference axis* appropriately draws the viewer’s attention to the magnitude and variability information in the two groups’ comparison (Cumming and Calin-Jageman, 2016). We can use this axis to diagram NHST (Figure 1D). This design has three main features. (1) The mean of the null is the difference-axis origin, zero. (2) The origin is flanked by a *sampling-error distribution*; this null distribution can constructed with permutation (Pitman, 1937). (3) The *P* value is visualized as the tail segment of the distribution that is more extreme than the observed effect size. If this tail segment is smaller than a predefined significance level, traditionally α = 0.05, an analyst will reject the null.

While visualizing the null distribution is an improvement, this picture nevertheless illustrates the flawed logic of NHST: in order to prove that the null hypothesis is false, the analyst must invoke the existence of something (the tail segment) that the hypothesis predicts (Berkson, 1942). NHST has been criticized for this fallacy, as well as its misleading dichotomization (McShane and Gal, 2017; Yildizoglu et al., 2015). Even the premise of NHST is incorrect: any intervention to any system will produce some (at least infinitesimal) effect, thus a hypothesis of a precisely zero effect size is inevitably false (Cohen, 1994).

### Estimation plots combine transparency and insight

As interventions always have effects, the analyst’s appropriate task is to quantify the effect size and assess its precision. First introduced by two biostatisticians over 30 years ago (Gardner and Altman, 1986), the best design for the analysis of two groups is an estimation plot that visualizes observed values and the effect size side-by-side. In this graphic, the difference-axis origin is aligned with the mean of the test group, making it easy to relate observed values to the *difference of means*, ∆ (Figure 1E). Around ∆, the analyst plots an indicator of precision known as the 95% *confidence interval* (CI) (Altman et al., 2000). The current design updates the Gardner-Altman plot by diagramming the sampling-error distribution as a filled curve.

### Five key advantages of estimation plots

Estimation plots possess five key advantages over conventional NHST plots. First, as mentioned above, the difference axis affords transparency of the comparison being made. Second, while *P* values conflate magnitude and precision in a single number, the relative size of a CI provides a specific measure of its precision. Third, plotting the full sampling-error curve of the effect size prevents dichotomous thinking and draws attention to the distribution’s graded nature. Fourth, deriving this sampling-error curve with bootstrapping makes the method robust and versatile. Fifth, and most importantly, by focusing attention on an effect size, the difference diagram encourages quantitative reasoning about the system under study. Such reasoning empowers scientists to make domain-specific judgements on whether an effect magnitude is noteworthy and relevant to their research question.

### Estimation plots are accessible

As two-groups analysis is the most frequently used statistical method in experimental research (Chochlac, 2018), the broad adoption of estimation for this type of analysis would greatly advance its acceptance more generally. However, while every major data-analysis tool can perform a Student’s *t*-test and chart NHST plots, very few software packages offer estimation plots. To improve the accessibility of estimation plots, we developed Data Analysis with Bootstrap-coupled ESTimation (DABEST), available in three open-source libraries for Matlab, Python, and R. DABEST calculates the sampling distribution and the CI with bootstrapping: resampling with replacement from the observations several thousand times (Efron, 1979; Efron and Tibshirani, 1994). Compared to parametric methods, bootstrapping is more robust for data sets with non-normal distributions (Efron, 1981, 1987). We have also used DABEST to build a free, user-friendly web application: estimationstats.com. Data is input via a spreadsheet, summary statistics are downloadable as text tables, and plots can be saved in image formats suitable for publication (PNG and SVG). The default CI can be easily re-specified to accommodate other interval sizes (Benjamin et al., 2017). With the web application and open-source libraries, DABEST caters to both scripting and spreadsheet workflows, empowering all researchers to rapidly adopt better data-analysis practices.

### Estimation plots are versatile

DABEST can be used to visualize large samples (Figure 2A), paired data (Figure 2B), multiple groups (Figure 2C), shared-control designs (Figure 2D), and to display standardized effect sizes such as Hedges’ g. More generally, estimation-focused plots can be used for linear regression (Figure 3), and meta-research (e.g. forest plots) (Borenstein et al., 2009). Thus, the estimation approach is broadly relevant (Claridge-Chang and Assam, 2016).

**Figure 2.**
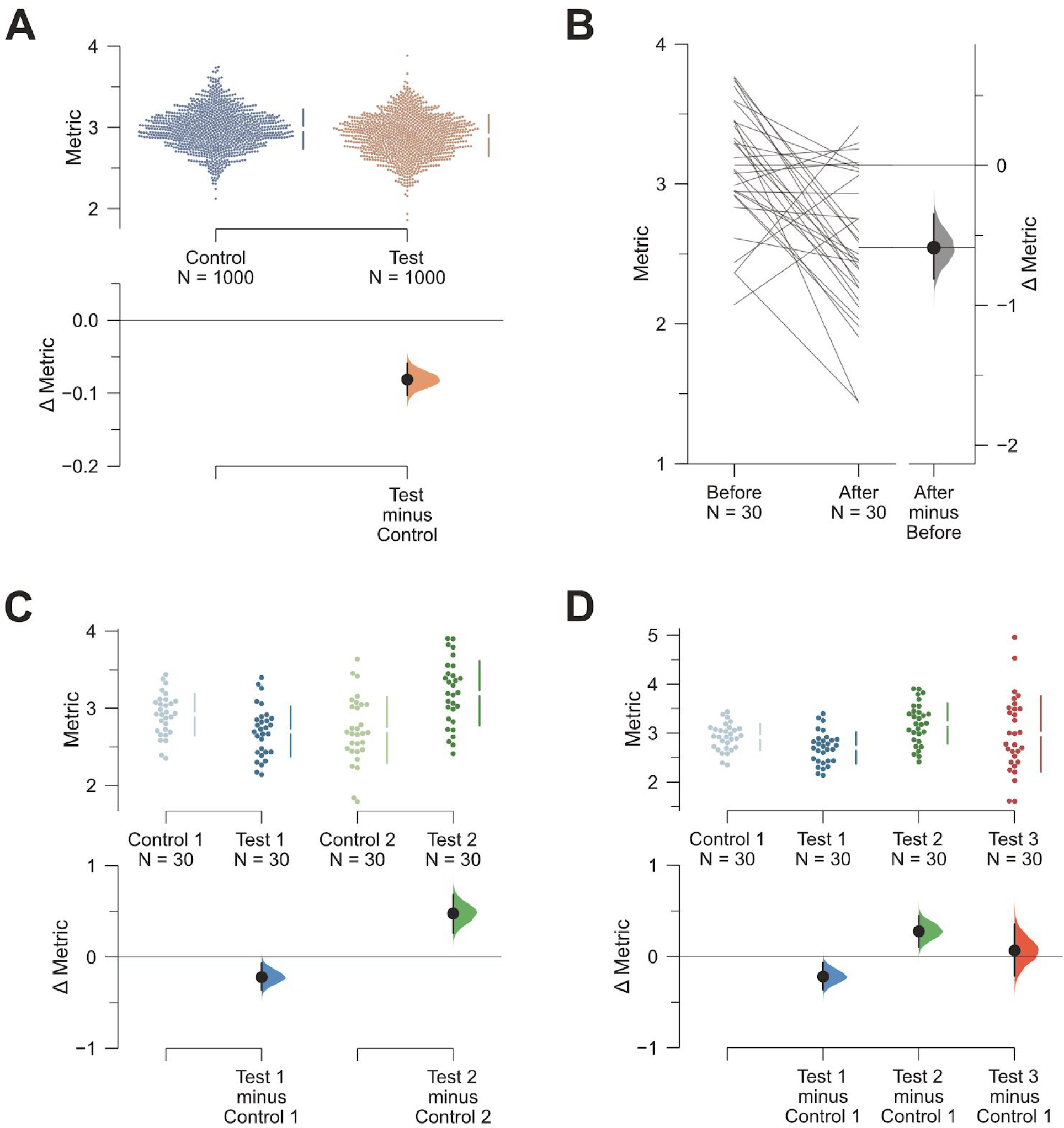
The estimation approach can accommodate a range of experimental designs. **A**. Estimation plots can effectively display datasets with large numbers of observations. Shown here is a plot of the mean differences between two groups, each with 1000 observations. The mean and standard deviation (SD) of each group is plotted as a gapped line (the ends of the vertical lines correspond to mean ± SD, while the mean itself is depicted as a gap in the line) alongside all data points. Even larger samples are easily handled with related designs: the violin plot (Hintze and Nelson, 1998) which shows the density of observed values; or the sinaplot (Sidiropoulos et al., 2017) which controls the jitter width of the data points according to each group’s density function. **B**. An estimation plot with a slopegraph (Tufte, 2001) depicting the pairs of within-subject observations and the ∆. **C**. A multi-group estimation plot with multiple two-group comparisons plotted together. The lower panel—which shows the effect sizes (Δs)—is analogous to a forest plot, summarising the results of several comparisons. We propose multi-group estimation plots be named ‘Cumming plots’ after their originator (Cumming, 2012). **D.** A shared-control plot. This is analogous to an ANOVA with multiple comparisons, where several groups (in this case, three groups: Test 1, Test 2, and Test 3) are compared against a single common control or reference group.

**Figure 3.**
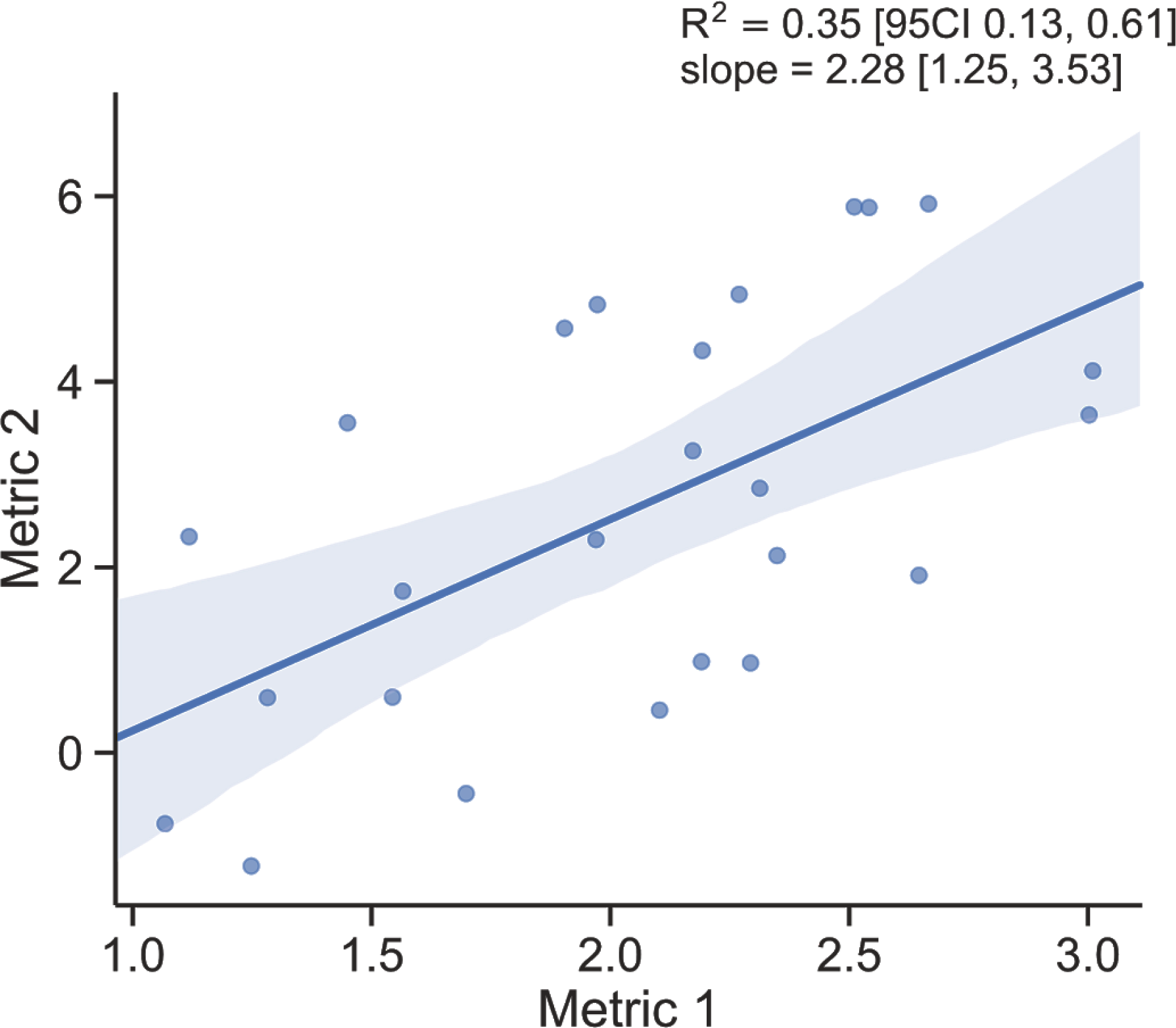
The estimation approach to linear regression. The principle of showing both observed values and effect size applies to other types of estimation plots. Shown here is the example of a linear regression plot. A best-practice visualization should include the following: (1) a scatter plot that shows all observations; (2) a fit line with its confidence-interval band; (3) the slope effect size (*m* in *y* = *mx* + *c* of the regression fit line) with its confidence interval; and (4) the coefficient of determination effect size (R^2^) with its confidence interval. These features of a regression plot are the counterparts to the key aspects of estimation graphics for grouped data: dot plot; difference axis; ∆ value; and standardized effect size (e.g. Cohen’s *d* or Hedges’ g) (Cohen, 1988; Hedges, 1981). Both *m* and ∆ express a change in the dependent variable in terms of the independent variable—continuous or categorical, respectively. Both R^2^ and *d* are indicators of the effect size as a proportion of variance; indeed, there are formulas for the interconversion of R^2^ and d-type effect sizes (Borenstein et al., 2009).

## Conclusion

Estimation plots constitute an elegant, robust framework for presenting data. The three software packages and accompanying web application offer non-statisticians a way to analyze their data without recourse to NHST, which derails analysts from quantification and misleads them to settle for superficial dichotomies. By visualizing effect sizes and their precision, estimation plots help analysts focus on quantitative thinking, enabling better scientific practice.

## Author Contributions

Conceptualization: JH, ACC; Methodology: JH, ACC; Software: JH (Python, R), TT (Matlab, R), SA (Matlab); Writing: Original Draft: JH, Revision: JH, HC, ACC; Visualization: JH, ACC; Supervision: HC, ACC; Project Administration: ACC; Funding Acquisition: HC, ACC.

## Acknowledgements

The authors are grateful to Mashiur Rahman for help and advice, and to Hung Nguyen for developing the web app front end.

## Sources of Funding

JH was supported by the A*STAR Scientific Scholars Fund. TT was supported by a Singapore International Graduate Award from the A*STAR Graduate Academy. SA was supported by a Singapore International Pre-Graduate Award. HC was supported by grants MOE-2016-T2-1-001 from the Singapore Ministry of Education and NMRC-CG-M009 from the National Medical Research Council. ACC was supported by grants MOE-2013-T2-2-054 and MOE2017-T2-1-089 from the Singapore Ministry of Education, grants 1231AFG030 and 1431AFG120 from the A*STAR Joint Council Office, and Duke-NUS Medical School. The authors received additional support from a Biomedical Research Council block grant to the Institute of Molecular and Cell Biology.

## Data Availability

The <u>Matlab</u>, <u>Python</u>, and <u>R</u> packages are all available on Github, and are licensed under the BSD 3-Clause Clear License.

## Guide to using DABEST

There are five ways to use DABEST.

No installation or download is required for the web application or Google Colab; either requires only an internet connection. The other methods require you to install Python, Matlab, or R on your personal computer.

## Web application

1. Access estimationstats.com.
2. Choose one of the functions, e.g. two groups.
3. Use the preloaded data or enter your own data.

## Google Colaboratory

1. Open an window in any modern browser (Chrome, Firefox, or Safari). Use *incognito* or *private* mode if you wish to remain anonymous.
2. Access this online example notebook to view the code that generated the Figure. You can view or download the notebook, but cannot run it without signing in.
3. If you would like to run the code in Colaboratory, you will need an Google account with which to sign in.

## Matlab

1. Download DABEST-Matlab from Mathworks File Exchange or the Github repo.
2. Follow the tutorial on Github.

## Python

1. Install the Anaconda distribution of Python 3.6 and Jupyter.
2. Download the example notebook from Colaboratory (see above).
3. Run the example notebook to install and test DABEST-Python.
4. Or, install DABEST with this line in the terminal:

~~~
pip install dabest
~~~
5. A tutorial on DABEST-Python can be found here.

## R

1. Run this line in the R console:

~~~
install.packages(“dabestr”)
~~~
2. Note that a version of R > 3.5.0 is required.

